# Opening of a cryptic pocket is not suppressed by excluded volume effects of crowding

**DOI:** 10.1101/2025.11.17.688865

**Authors:** Prajna Mishra, Si Zhang, Gregory R. Bowman

## Abstract

Cryptic pockets are transient structural features that could provide new opportunities for drug design. However, their utility would be greatly diminished if the probability of pocket opening were universally suppressed in cells due to the excluded volume effects of crowding. As a first step to addressing this concern, we study the effect of the synthetic crowder Ficoll 70, which has been shown to mimic the excluded volume effects of biomolecular crowding well, on the opening of a cryptic pocket in the interferon inhibitory domain (IID) of Ebola viral protein 35 (VP35). We find that this synthetic crowder has a negligible effect on the probability of cryptic pocket opening, as measured by both thiol labeling and hydrogen-deuterium exchange mass spectrometry (HDX-MS). The probability that a specific pocket opens may still be altered by the soft, enthalpic interactions of crowding, which will vary between different cellular environments. However, our results demonstrate that excluded volume effects that are at play in all crowded environments do not pose a universal barrier to targeting cryptic pockets.

## Introduction

Cryptic pockets are sites that open in excited states but are closed in the ground state and, therefore, are typically absent in experimentally derived protein structures.^1, 2^ They have rapidly gained attention as drug targets for challenging protein-protein or protein-nucleic acid interactions, particularly for proteins considered "undruggable" due to the lack of apparent binding grooves in their ground-state structures.^3, 4^ Because they are often allosterically coupled to functional regions, cryptic pockets provide unique opportunities to modulate protein activity in ways inaccessible to conventional active-site focussed drug design.^5^ Beyond broadening target space, cryptic pockets can enable selective modulation; they may discriminate between isoforms or homologs, and their allosteric nature allows both inhibition and enhancement of protein function, in contrast to the primarily inhibitory outcomes of traditional active-site ligands.^5-7^ Recent large-scale analyses suggest that at least half of proteins lacking a canonical binding groove may, however, harbor cryptic sites, greatly expanding the druggable proteome across diverse biological processes, including immunity, signaling, and cancer.^8^

While much has been learned about cryptic pockets from biophysical studies performed in dilute conditions,^9^ it is unclear whether they behave the same in crowded cellular environments. In living cells, proteins and nucleic acids exist in a highly crowded environment, where macromolecules can occupy between 8 - 40% of the cytoplasmic volume.^10-12^ Such macromolecular crowding can significantly influence protein folding, stability, and conformational equilibria.^13-20^ Crowding’s effects are often decomposed into two components. The first is excluded volume effects, which are present in every crowded environment and depend mainly on the volume occupied.^21-23^ The second component is soft, enthalpic interactions (called quinary interactions) between a protein of interest and the macromolecules crowding it.^23-25^ The effects of these interactions are highly dependent on the chemistry of the specific crowder(s) and, therefore, vary between cellular contexts.^21, 24^

A first question with cryptic pockets is whether the excluded volume effects that are at play in all crowded conditions have an impact on the probability of pocket opening. These excluded volume effects are generally thought to stabilize proteins by disfavoring extended or partially unfolded conformations.^19-22^ Because cryptic pockets often require local expansion of the protein surface, pocket opening could be suppressed under crowded conditions. If such effects reduced the probability of cryptic pocket opening, then that would likely mean that many cryptic pockets would not be viable drug targets. If any cryptic pocket is not suppressed by excluded volume effects, then one can conclude that excluded volume effects of crowding do not universally suppress pocket opening. Of course, quinary interactions within a specific environment could still mean that some environments suppress pocket opening.

In this study, we use the synthetic crowder Ficoll 70 to test if excluded volume effects universally suppress cryptic pocket opening. Synthetic crowding agents such as dextran, Ficoll, and polyethylene glycol (PEG) have been widely used to study excluded volume effects.^13, 26^ Of these, Ficoll 70 is particularly good at mimicking the excluded volume effects of macromolecular crowding, as its branched, quasi-spherical structure makes it more similar to globular proteins than more flexible polymers like PEG are.^13^ Combining it with other reagents meant to mimic aspects of crowding has even yielded promising cytomimetics. For example, studies on phosphoglycerate kinase (PGK) and the bacterial surface protein VlsE showed that a mixture of ∼15% w/v Ficoll 70 and lysis buffer recapitulated cell-like stability and kinetics.^27^ Similarly, conditions containing ∼15% w/v Ficoll 70 supplemented with 20% v/v mammalian cell extraction reagent (M- PER™) have been shown to provide a broadly applicable cytomimetic environment for λ-repressor protein fragment 6–85 (λ_6-85_).^15^ Based on these established precedents, we go up to 40% w/v Ficoll 70 to mimic the excluded volume effects of the maximum crowder concentration reported in cells.^10-12^

As a model system, we chose a cryptic pocket in the C-terminal interferon inhibitory domain (IID) of Ebola virus protein 35 (VP35). We will refer to this single domain as VP35 for brevity. This protein binds to double-stranded RNA (dsRNA) and blocks RIG-I-mediated signaling, a critical step in immune evasion. Even modest disruption of this interaction impairs viral fitness, highlighting VP35 IID as a promising therapeutic target.^28-30^ Previous work from our group identified this cryptic pocket using molecular dynamics (MD) simulations (Figure 1A) and subsequently confirmed its existence experimentally through thiol labeling and dsRNA binding experiments.^5, 31^ All three techniques (MD, thiol labeling, and dsRNA binding) reported similar open-state populations (Figure 1B).^5^ The pocket forms when the helix spanning residues 303–310 separates from the 4-helix bundle (Figure 1 A). The G236–A306 Cα–Cα distance cleanly distinguishes open and closed conformations (Figure 1C). These conformational changes are accompanied by minimal differences in helical propensity within the pocket region, indicating that pocket formation predominantly arises from coordinated helical rearrangement rather than local unfolding (Figure 1D). However, the helix can also unfold, leading to more extended open conformations.

**Figure 1:**
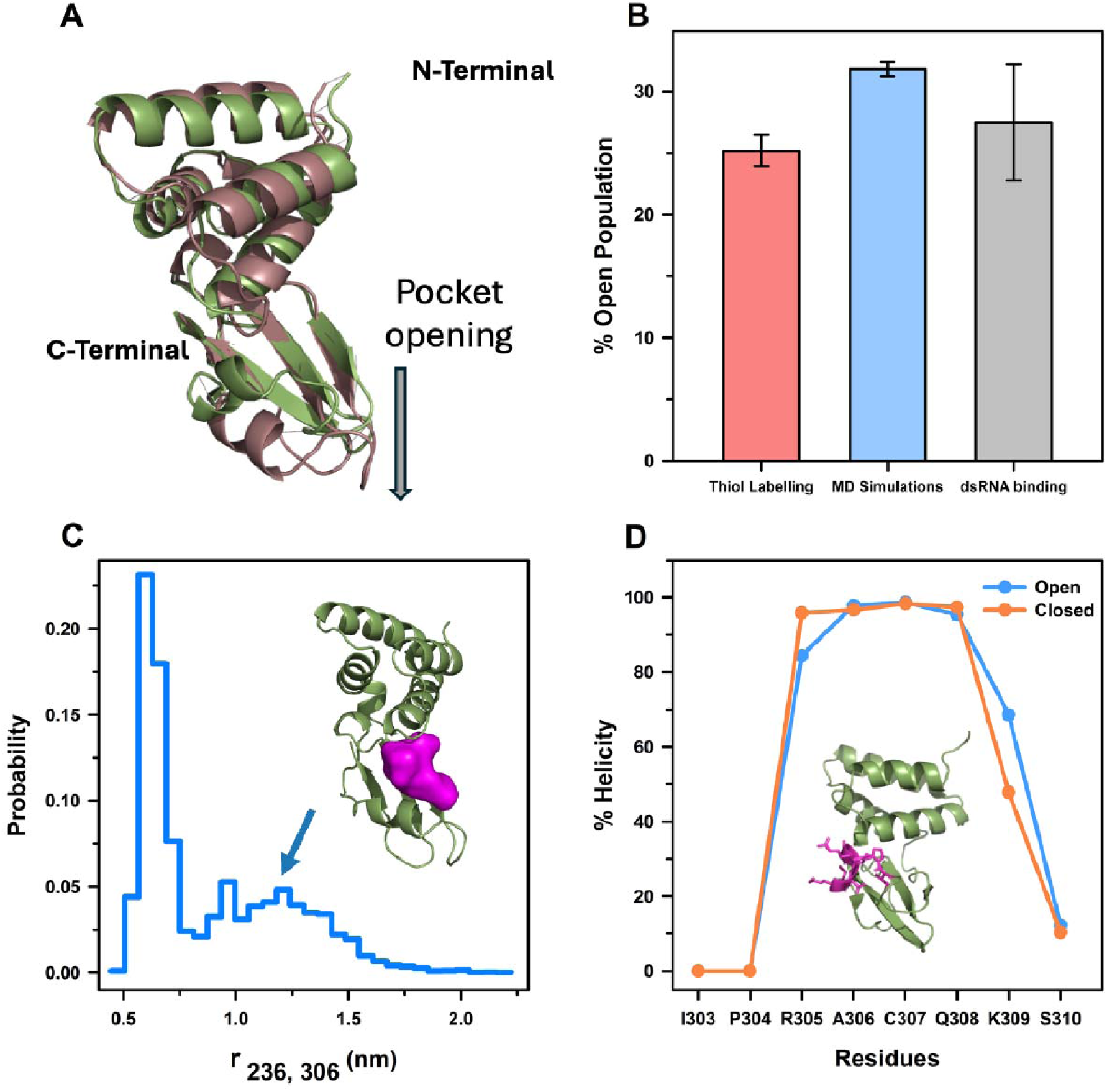
A cryptic pocket in Ebola VP35 IID captured by complementary computational and experimental approaches. (A) Structural overlay of VP35 IID in the closed (green) and open (brown) conformations. The conformational rearrangement exposes a cryptic pocket that is allosterically coupled to the dsRNA binding interface. (B) Open state populations estimated using independent approaches. Bar plots show the percentage of pocket opening derived from thiol labeling experiments, molecular dynamics (MD) simulations and dsRNA binding experiments.^5^ Experimental uncertainties are reported as the standard deviation of triplicate measurements; computational uncertainties were estimated using a per-trajectory bootstrap procedure. (C) Distribution of the G236-A306 Cα-Cα distance from MD simulations. The inset shows the pocket volume (pink) calculated using LIGSITE for a representative open-state structure. There is plenty of room to accommodate a small molecule or even a peptide, and previous application of FTMap^32^ to our structures shows there are plenty of favorable interactions that drug-like molecules can form in this pocket.^31^ (D) Residue-level helical propensity comparison between open and closed ensembles derived from MD simulations. Percent helicity is plotted for residues spanning the pocket region (303-310). The inset highlights pocket residues 303-310 shown as magenta sticks. The uncertainty is negligible and is not shown for visual clarity.

We quantify the probability of pocket opening in the presence of different levels of Ficoll 70 using thiol-labelling kinetics and hydrogen-deuterium exchange mass spectrometry (HDX-MS). Thiol labeling reports on the solvent exposure of a cysteine sidechain within a cryptic pocket and is particularly well-suited to pockets where the open state predominantly maintains secondary structure, as we have seen for VP35. Indeed, we have previously established that the labeling rate of C307 reports on the probability that our VP35 cryptic pocket is open.^5^ HDX-MS reports on the solvent exposure of backbone amides. Therefore, we expect it to report on the small fraction of the open population that is more disordered and, therefore, exposes backbone amides to solvent. More extended conformations tend to be more sensitive to the excluded volume effects of crowding, so HDX-MS experiments are a particularly sensitive reporter on any effects that excluded volume has on our model cryptic pocket.

## Results

### 1. Thiol labeling reveals that the VP35 cryptic pocket is not suppressed by excluded volume effects

To quantify the impact of the excluded volume effects of crowding on the probability of pocket opening, we first turned to thiol labelling experiments. These experiments are particularly suitable probes in this context because they report on side-chain degrees of freedom, which may change even when secondary structure is preserved.^31, 33, 34^ This feature is an important advantage since our simulations suggest more pronounced changes in sidechain exposure than in the backbone degrees of freedom that are detected by HDX-MS (Figure 1D). Using a drug-sized labeling reagent also provides insight into whether the pocket opens enough to accommodate a small molecule drug. Of the four native cysteines in the VP35 IID, two (C307 and C326) are located within the cryptic pocket, one (C275) maps to the dsRNA-binding interface, and one (C247) is buried in the helical bundle (Figure 2A). Prior work from our group has shown that thiol exchange effectively tracks site-specific cysteine accessibility in this domain.^5, 31^Among these, C307 is the most informative reporter of pocket opening. Its solvent exposure correlates strongly with pocket dynamics and has been linked to modulations of dsRNA binding.^5, 31^ Building on these observations, we focused on C307 as a sensitive probe of cryptic pocket accessibility under crowded conditions.^5^ We hypothesized that if the excluded volume effects of crowding favor the closed state of the pocket, the accessibility of C307 would decrease; conversely, unchanged accessibility would suggest that the pocket remains dynamic in crowded environments. We chose to focus on Ficoll 70 as it has been shown to mimic the excluded volume effects of cellular environment^15, 27^ and it behaves well in all the assays employed in this study.

**Figure 2:**
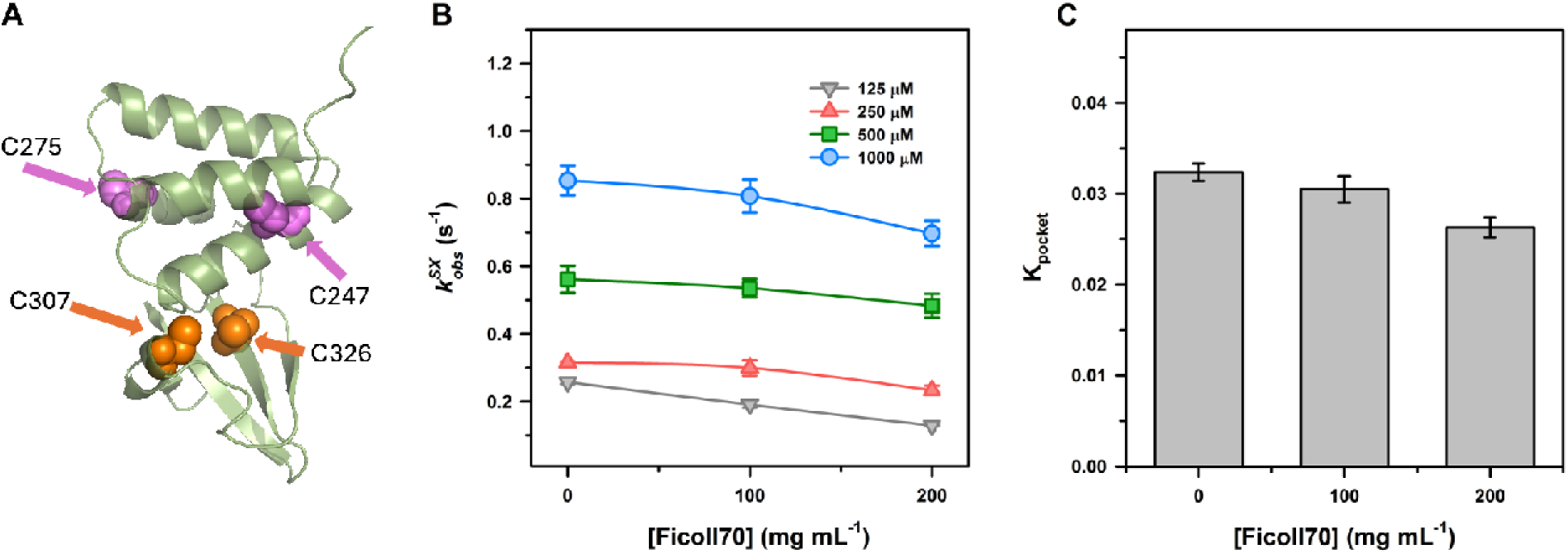
Cryptic pocket cysteine remains solvent accessible under crowding by Ficoll 70. (A) Structure of the VP35 IID highlighting four native cysteines: cryptic pocket cysteine C307 (orange sphere), C326 (orange sphere), C275 on the dsRNA binding interface (purple sphere), and buried C247 in the helical bundle (purple sphere). (B) Observed labeling rates () for C307 in VP35 C247S/C275S variant, measured across Ficoll 70 concentrations and multiple DTNB concentrations (125-1000 μM). (C) Equilibrium constants of pocket opening (K_pocket_) across Ficoll 70 concentrations. Error bars in the graphs represent standard deviations from at least three independent measurements.

For these experiments, we used a C247S/C275S variant that only has cysteines in the cryptic pocket, which we have previously shown allows us to study the pocket without the confounding effects of labeling cysteines elsewhere in the protein. Thiol exchange was carried out using 5,5′- dithiobis (2-nitrobenzoic acid) (DTNB), which reacts with exposed thiols to release TNB, resulting in a measurable absorbance increase at 412 nm (Supplementary Figure 1). Pocket opening events are thus detected as exponential increases in absorbance corresponding to cysteine modification. Labeling was measured across multiple DTNB concentrations (125-1000 μM) and Ficoll 70 concentrations (0-200 mg/mL) to obtain kinetic parameters. The equilibrium probability the pocket is open and the opening rate were inferred by fitting data at multiple DTNB concentrations with the Linderstrøm-Lang model, as described in Materials & Methods. Ideally, experiments would have been performed up to 40% w/v Ficoll 70 in buffer, approximating the highest macromolecular crowding levels reported in cells. However, above 200 mg/mL of Ficoll 70 the viscosity slows the diffusion of DTNB so much that one cannot disentangle the viscosity effects from the behavior of the pocket. Even so, the 20% w/v we are able to make measurements for is well above the 15% w/v Ficoll 70 that has been reported to mimic cellular environments in multiple studies.^15, 27^

We observed that C307 remained solvent accessible under crowded conditions, demonstrating that the cryptic pocket persists as a dynamic feature even in the presence of crowders. In a dilute buffer, C307 was solvent accessible, showing robust labeling consistent with dynamic pocket opening (Figure 2B). Importantly, increasing Ficoll concentrations up to 20% w/v had little effect on the observed labeling rate 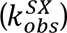 and equilibrium constants of pocket opening (K_pocket_) as shown in Figures 2B and 2C, respectively. There is some reduction in the equilibrium constant at 20% w/v but the change is not large, and we suspect it is likely due to issues arising from the solvent viscosity rather than a real change in the protein’s behavior (Figure 2C, Supplementary Figure 2). This result is good evidence that the excluded volume effects of crowding do not suppress cryptic pocket opening given that ∼15% w/v Ficoll 70 has been found to mimic cellular crowding.^15, 27^ It would be even stronger to go up to 40% w/v. However, higher Ficoll levels could not be tested due to excessive viscosity, which interfered with mixing. Fitting the DTNB-concentration dependence yielded pocket-opening rates of 1.60 s^- 1^, 1.68 s^-1^ and 1.66 s^-1^ at 0, 100, and 200 mg/mL Ficoll 70, respectively, with no systematic decrease under crowded conditions (Supplementary Figure 3). The fact that DTNB is roughly drug-sized also demonstrates that the pocket is opening enough to accommodate a small molecule drug across these conditions. The thermal stability of the protein also remains constant even at 40% w/v of Ficoll 70 (Supplementary Figure 4). Together, our thiol labeling results demonstrate that pocket opening is largely unaffected by the excluded volume effects of crowding up to 20% w/v and the global stability data suggests that the trend may hold up to 40% w/v, though a more quantitative readout at these higher crowding concentrations would b stronger.

### 2. HDX-MS captures cryptic pocket opening

We next turned to HDX-MS to reach higher Ficoll 70 concentrations that better match the most extreme crowding observed in cells. Unlike thiol exchange, HDX-MS reports on backbone amide exchange rather than side-chain reactivity, and its kinetics are governed by local hydrogen bonding and conformational dynamics.^35-38^ This makes HDX particularly well suited to probing protein dynamics in crowded environments. In our thiol labeling experiments, increased solvent viscosity due to crowding can slow mixing, making it hard to determine if slower labeling is due to pocket closure or slower diffusion of the chemical probe. In HDX, the chemical probe is the solvent, which is at such high concentration that slowed diffusion isn’t a concern. However, our simulations suggest that secondary structure is largely the same whether the pocket is open or closed (Figure 1D). This could present a challenge for HDX-MS as it reports on exchange of the backbone amide, which is suppressed when secondary structure is formed. Therefore, HDX-MS could report on the subset of the ensemble with an open cryptic pocket where secondary structure is lost, or it could be insensitive to pocket opening. Before performing HDX-MS under different crowding conditions, we first set out to test if it detects enhanced exchange in the cryptic pocket compared to other parts of the protein.

We performed global HDX-MS on wild-type (WT) VP35 IID to determine whether the cryptic pocket could be identified from backbone exchange profiles. An overlapping peptide map of the WT VP35 IID was generated by pepsin digestion, providing 100% sequence coverage with an average redundancy of 3-5 peptides per residue (Figure 3A, Supplementary Figure 5). Samples were incubated in D O-containing buffer for time points ranging from 20 s to 24 h, after which exchange was quenched, peptides were separated by liquid chromatography, and deuterium incorporation was quantified by MS. Percent deuterium uptake was then calculated for each peptide as a function of labeling time, producing baseline uptake profiles across the IID (Figure 3).

**Figure 3:**
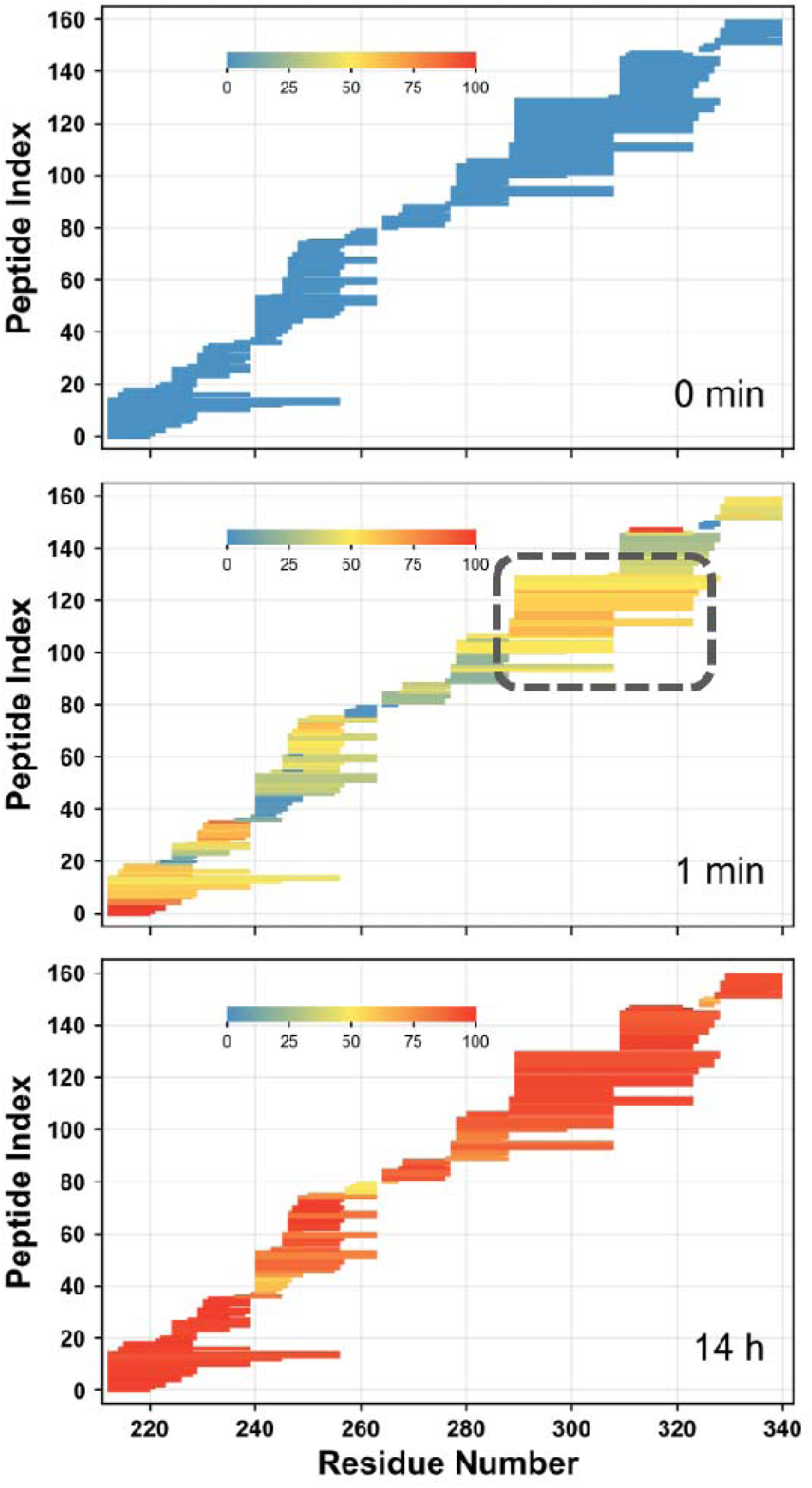
Global HDX-MS delineates the cryptic pocket in VP35 IID at peptide resolution. (A-C) Percent deuterium uptake (%D) for overlapping peptides plotted by sequence position (x- axis, residue number 211-340) and peptide index (y-axis). Color encodes uptake from 0% (blue) to 100% (red). (A) 0-min control (back exchange baseline) confirms broad peptide coverage with minimal initial uptake. (B) At 1 min, the cryptic pocket region (dashed box) shows higher deuterium uptake than the rest of VP35 IID, exchanging more than 60%. (C) 14-h labeling approaches equilibrium uptake. Bars represent unique overlapping peptides; the data shown are representative of at least triplicate measurements.

At 1 min, more than 60% of amide hydrogens within peptides spanning the cryptic pocket region had exchanged, marking this segment as highly dynamic relative to neighboring structured elements, which remained substantially more protected (Figure 3B). By 14 h, deuterium uptake approached saturation across nearly all residues (Figure 3C). These observations reveal that the cryptic pocket resides in a locally flexible segment that transiently fluctuates from the rigid helical/β sheet scaffold. The results also show enhanced exchange in the first helix of the protein. This observation agrees with our previous work, wherein we saw displacement of the first helix from the helical bundle in our simulations and observed labeling of C247, which is buried in available crystal structures but becomes exposed when the N-terminal helix is displaced.^31^ The exchange behavior observed here is consistent with EX2 exchange, in which observed HDX rates report on the equilibrium population of exchange-competent conformations rather than on separate open and closed states, and resulting in continuous centroid shifts rather than bimodal isotopic envelopes. Importantly, the HDX signal cannot be explained by global unfolding. Based on the measured global stability of VP35 IID (5.69 ± 0.35 kcal/mol), the fully unfolded population is only ∼0.007%, far below the exchange-competent population inferred from HDX-MS (∼0.03% for the major component). Therefore, HDX-MS is most consistent with local pocket-region fluctuations rather than global unfolding. This interpretation is also consistent with our model of the VP35 cryptic pocket: thiol labeling detects side-chain exposure of C307 in the dominant open-pocket ensemble, whereas HDX-MS requires backbone-amide exposure and therefore reports on a smaller, more expanded (e.g. locally disordered) subset of open conformations. Because these expanded conformations should be especially sensitive to excluded-volume effects, any crowding-induced suppression of pocket opening should be readily apparent as increased protection in the pocket-spanning peptides. Together, these results demonstrate that HDX-MS provides a complementary readout of the VP35 IID cryptic pocket region and establishes a baseline for evaluating the effects of macromolecular crowding on its dynamics.

### 3. HDX–MS reveals that the probability of cryptic pocket opening is unchanged by 40% w/v Ficoll 70

Having established that HDX-MS delineates the cryptic pocket as a highly dynamic segment of VP35 IID, we next asked whether these dynamics are sensitive to the excluded volume effects of crowding. Specifically, we tested whether the cryptic pocket is stabilized under extreme crowding by using 400 mg/mL (40% w/v) Ficoll 70 in buffer, approximating the highest macromolecular crowding levels reported in cells.^10, 11^

We measured the kinetics of deuterium incorporation into multiple overlapping peptide fragments spanning the cryptic pocket, with labeling times up to 24 h (Supplementary Figure 5). Percent deuterium uptake for each peptide, corrected for back-exchange, was fitted to exponential models (Supplementary Figure 6). From these fits, the observed exchange rates 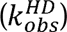 were obtained. The observed rates were then normalized to the intrinsic exchange rates (*k_int_*) to calculate the protection factor (PF) for each peptide. Multiple overlapping peptides spanning the pocket region showed similar protection factors, indicating that they report on a shared local opening process rather than global unfolding. Thus, the HDX-MS signal from pocket peptides reflects local pocket-region fluctuations rather than global unfolding.

As expected, percent deuterium (%D) uptake for cryptic pocket peptides remained nearly identical from zero crowder to the highest concentration (40% w/v), indicating the probability of pocket opening is constant. Representative peptides D289-Q308 and K309-F328, spanning the cryptic pocket and the adjacent exposed region, are shown in Figures 4A and 4B, respectively. Both peptides exhibited similar deuterium uptake profiles in buffer versus 400 mg/mL Ficoll 70, approximating the highest macromolecular crowding reported in cells. This invariance demonstrates that the microenvironmental hydrogen bond network in the pocket region is unaffected by the excluded volume effects of crowding, implying that the relative probability of open and closed conformers is preserved. Under EX2 exchange conditions, protection factors 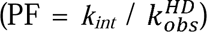 are inversely proportional to the equilibrium constant for local unfolding or solvent exposure.^39^ Thus, unchanged PF values indicate that the equilibrium constant for pocket opening (*K_op_*) is unaltered by the excluded volume effects of crowding. For peptide D289–Q308, protection factors calculated from the three kinetic phases were comparable under both conditions. For example, the dominant phase (61.46 ± 3.2% amplitude) had a protection factor of 2.84 10 0.47 10 in buffer and 8.96 10 1.64 10 in Ficoll (60.76 ± 2.72% amplitude) (Table 1), corresponding to open-state populations of approximately 0.03% and 0.01%, respectively. For K309-F328, the dominant phases likewise showed overlapping protection in buffer versus in Ficoll. These quantitative comparisons confirm that pocket peptides retain nearly identical backbone protection in crowded versus dilute solution. Collectively, these data demonstrate that even under 40% Ficoll 70, approximating maximal cellular crowding, the VP35 IID cryptic pocket dynamics remain unaltered. Thus, the VP35 IID cryptic pocket provides an experimentally validated case in which functionally relevant hidden conformations remain populated even under macromolecular densities that mimic physiological conditions. The HDX-MS data show that even highly exposed backbone-exchange behavior in the pocket region is not suppressed at 40% Ficoll 70. Therefore, it is unlikely that the less globally extended, DTNB-accessible conformations detected by thiol labeling would be selectively suppressed only at higher Ficoll 70 concentrations. Indeed, that is what we see in our thiol labeling experiments going up to 20% w/v of crowder. For our conclusion to be wrong, there would have to be some non-obvious physical factor that came into play at crowder concentrations from 20-40% w/v but not at lower concentrations, and that factor would have to somehow suppress opening of helical conformations while having negligible impact on the probability of more extended conformations.

**Figure 4:**
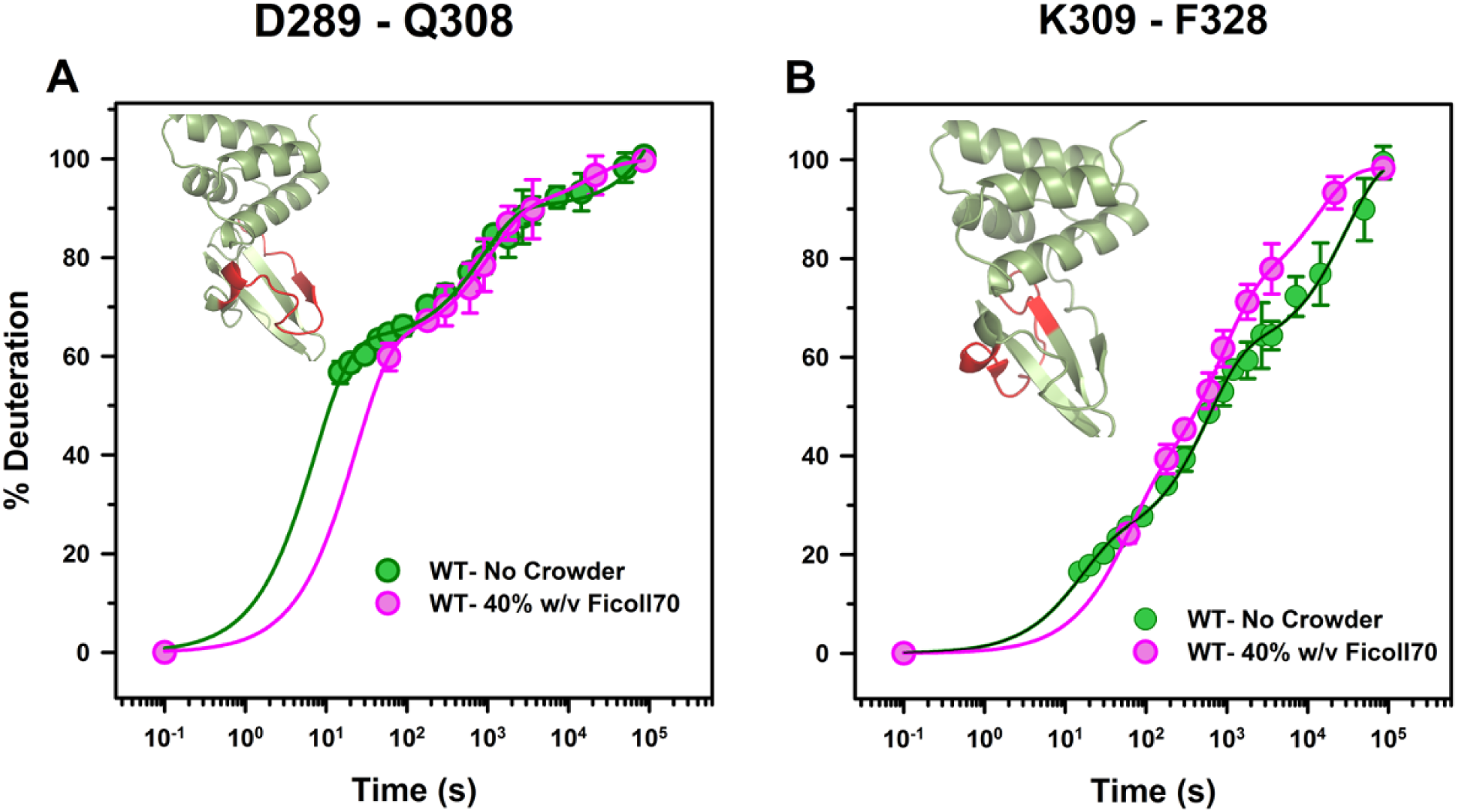
Representative deuterium uptake curves for cryptic pocket peptides show consistent dynamics even at 400 mg/mL of Ficoll 70, mimicking maximal crowding reported in cellular environments. (A) Uptake kinetics for peptide D289-Q308 in wild-type VP35 IID in buffer (green) and with 40% w/v Ficoll 70 (pink). (B) Uptake kinetics for peptide K309-F328 under the same conditions. Inset cartoons show the VP35 IID structure with the respective peptide highlighted in red. Data points represent mean and standard deviations in % deuterium uptake from at least three independent measurements; lines show best-fit exchange kinetics.

**Table 1.**
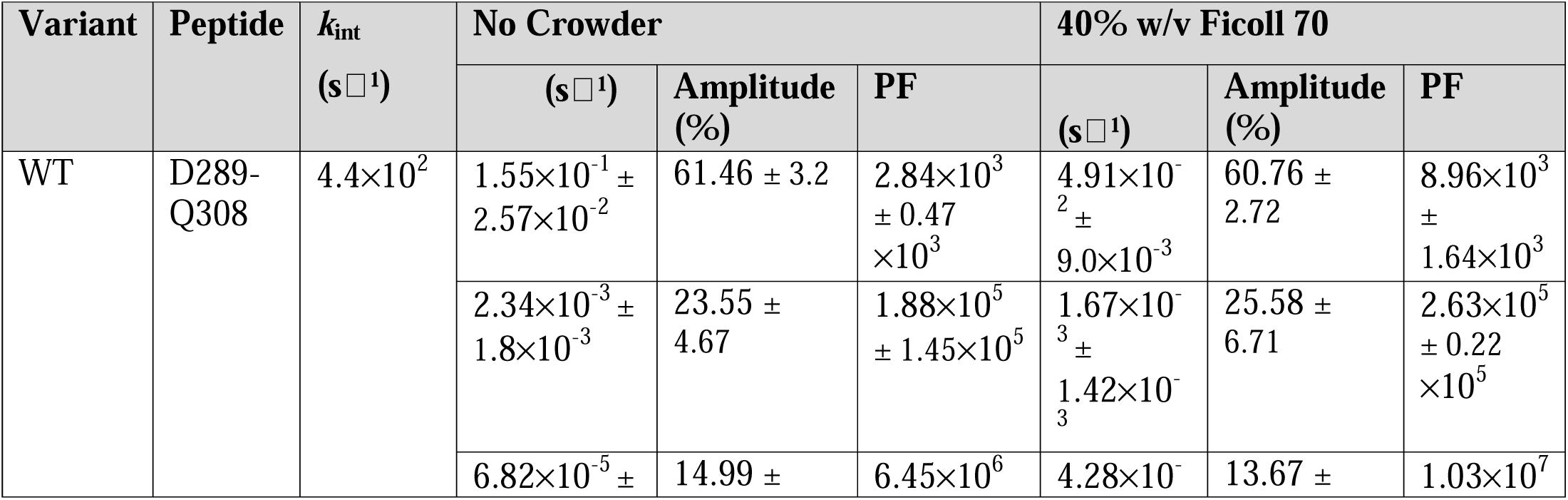
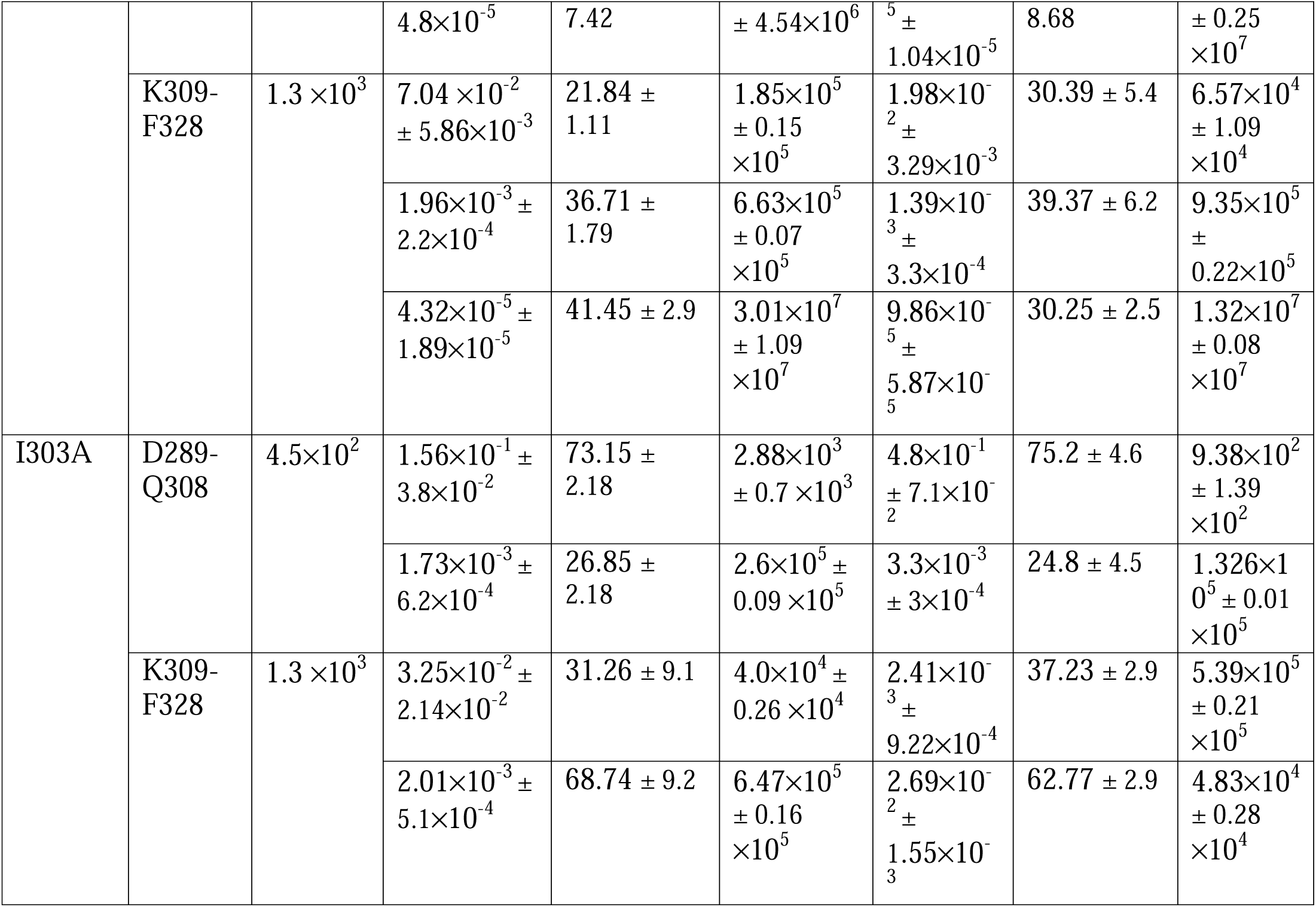
Observed exchange rate constants 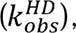 amplitudes, and calculated protection factors (PF) for cryptic pocket peptides in WT and I303A VP35 IID under buffer (no crowder) and 40% w/v Ficoll 70 conditions. PF was calculated as 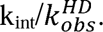 Values are shown to two significant figures.

### 4. Excluded volume effects do not suppress mutation-enhanced pocket opening

We reasoned that VP35 variants with a higher probability of cryptic pocket opening could reveal if the excluded volume effects of crowding have subtle effects that aren’t apparent for the wild-type protein. A small percent change on an already small population may go unmissed, whereas the same percent change for a larger population may be visible. To probe whether crowding could influence a pocket that samples the open state more frequently, we turned to PocketMiner, a graph neural network developed in our lab that predicts cryptic pocket locations and opening probabilities from static structures,^8^ to identify mutations that increase the probability of pocket opening in VP35. Screening all point mutations of VP35 suggested that the substitution I303A should enhance the probability of pocket opening, and thiol exchange experiments (Supplementary Figure 7) confirmed this prediction. Specifically, the probability of opening in WT VP35 IID is 25.19% ± 1.29% while the probability of opening in I303A is 65.71% ± 0.38%. We therefore asked whether the excluded volume effects of crowding could counteract these enhanced dynamics by stabilizing the closed state, or whether the mutation’s effect would persist even under crowded conditions.

HDX-MS was performed on I303A VP35 IID under the same conditions as WT VP35: in buffer and in 40% w/v Ficoll 70, approximating maximal cellular crowding. Deuterium uptake was monitored for representative cryptic pocket peptides D289-Q308 and K309-F328. Deuterium uptake kinetics were fitted to triple exponential models to obtain observed exchange rates 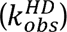 and corresponding PF (Table 1), enabling direct comparison of pocket dynamics between WT and I303A in dilute and crowded environments.

The I303A mutant behaved as expected: the cryptic pocket was more open than in WT VP35, and this enhanced opening was unaffected by the excluded volume effects of crowding. In buffer, I303A showed higher deuterium uptake for pocket peptides D289-Q308 and K309-F328 compared to WT, consistent with enhanced pocket opening (Figure 5A-B). This result also corroborates our claim that HDX-MS is detecting pocket opening, since the mutation impacts the cryptic pocket but produced minimal changes in deuterium uptake outside the pocket region (Supplementary Figure 8). Quantitative fitting confirmed reduced protection in I303A (Table 1). Unlike WT, whose exchange kinetics were best described by three exponential phases, the HDX profiles of I303A were adequately described by a two-phase model, consistent with a shift toward fewer, less-protected conformational states. Importantly, these differences persisted in 40% Ficoll 70: the uptake profiles of both peptides were nearly identical in buffer and Ficoll (Figure 5C-D, Supplementary Figure 9), and protection factors remained within the same range (Table 1). Together, these results demonstrate that Ficoll 70 does not suppress the destabilizing effect of the I303A mutation and that cryptic pocket dynamics remain conserved under crowded conditions even when the conformational equilibrium is shifted toward the open state.

**Figure 5:**
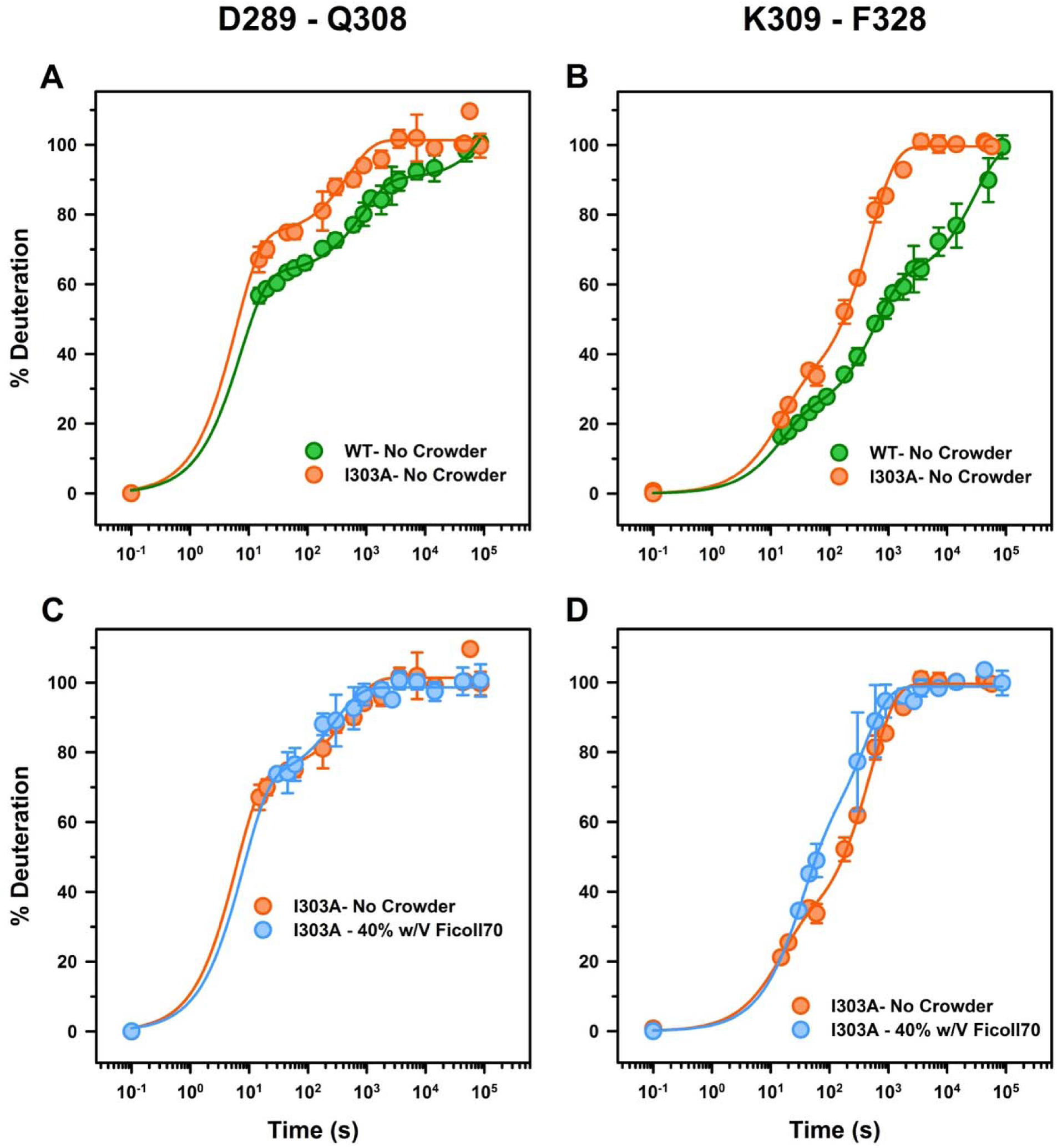
The I303A mutation promotes pocket opening compared to WT VP35; however, its effect on local dynamics remains conserved under Ficoll 70 crowding. (A-B) Deuterium uptake kinetics for pocket peptides D289-Q308 (A) and K309-F328 (B) comparing wild-type VP35 IID (green) and I303A mutant (orange) in buffer. (C-D) Deuterium uptake kinetics for the same peptides in the I303A mutant under buffer (orange) and 40% w/v Ficoll 70 (blue). Data points represent mean and standard deviations in % deuterium uptake from at least three independent measurements; lines show best-fit exchange kinetics.

## Discussion and Conclusion

Our results show that the probability of cryptic pocket opening in Ebola VP35 remains constant even with excluded volume effects under extreme macromolecular crowding (40% Ficoll 70). Across complementary experimental readouts, we did not observe substantial suppression of pocket opening. Thiol-labeling experiments showed that C307 remains accessible to DTNB under Ficoll 70 crowding, indicating that the pocket continues to sample conformations large enough to admit a ligand-sized small-molecule probe. HDX-MS further showed that backbone dynamics of peptides spanning the cryptic pocket are largely unchanged between dilute buffer and 40% w/v Ficoll 70, a condition approximating the upper range of macromolecular crowding reported in cells. Together, these data indicate that excluded volume effects do not suppress the VP35 cryptic pocket.

This conclusion is strengthened by experiments with variants that shift the pocket-opening equilibrium. The I303A variant, which strongly enhances pocket opening, has an open-state population of 65.71% ± 0.38%. Like the wild-type protein, this variant’s behavior was invariant up to 40% w/v Ficoll 70, demonstrating that excluded volume effects do not counteract mutation-enhanced opening or force the protein back toward the WT-like closed ensemble. Thus, the persistence of pocket opening under crowding is not limited to the lower open-state population of WT VP35 IID, but also applies to a variant in which the open state is substantially more populated.

We propose our conclusions are valid even though it was not possible to obtain thiol labeling data at crowder concentrations above 20% w/v. First, the relevance of our thiol labeling measurements up to 20% w/v is supported by the observation that 15% w/v of Ficoll 70 mimics multiple cellular environments.^15, 27^ Furthermore, our complementary measurements all suggest the probability of pocket opening is maintained up to 40% w/v. Our HDX experiments only report on the most extended conformations of the open cryptic pocket, where the backbone hydrogen bonds in the helical portion are broken. For our conclusions to be wrong, one would have to imagine a situation where crowding doesn’t impact the population of the most extended conformations of the open ensemble detected by HDX up to 40% w/v, doesn’t impact the population of less extended conformations detected by thiol labeling up to 20% w/v, but somehow has a significant effect on those less extended conformations in the range of 20-40% w/v.

Although our results demonstrate that excluded volume effects due to crowding do not suppress VP35 IID cryptic pocket opening, we cannot exclude the possibility that quinary (weak, nonspecific biomolecular) interactions modulate this equilibrium in specific cellular contexts. Extending these measurements to protein-based or mixed crowders represents a potential direction for future work, although implementing such systems in thiol-labeling and HDX-MS experiments presents substantial technical challenges.

## Materials and Methods

### Protein expression and purification

WT VP35 IID and all the variants were expressed in *E. coli* BL21(DE3) Gold cells (Agilent Technologies). Cultures were grown at 37 °C to an OD_600_ of 0.3, shifted to 18 °C, and further grown to an OD_600_ of 0.6 before induction with 1 mM IPTG (Gold Biotechnology). Following 24 h expression, cells were harvested and resuspended in lysis buffer (20 mM sodium phosphate, pH 8.0, 1 M NaCl, 5 mM β-mercaptoethanol).

Cells were lysed by sonication, and insoluble material was removed by centrifugation at 4 °C. The soluble fraction was purified by Ni-NTA affinity chromatography, followed by removal of the His-tag with TEV protease. Proteins were further purified by cation-exchange chromatography and finally by size-exclusion chromatography (Superdex 75). Purified proteins were stored in buffer containing 10 mM HEPES (pH 7.0), 150 mM NaCl, 1 mM MgCl2, and 2 mM TCEP.

Protein concentration was determined by absorbance at 280 nm using a calculated extinction coefficient of 6970 M^-1^cm^-1^. Purity and monodispersity were confirmed by SDS-PAGE and intact MS.

### Macromolecular crowding conditions

To mimic intracellular crowding, purified VP35 IID was incubated in storage buffer supplemented with various concentrations of Ficoll 70. These concentrations approximate physiologically relevant macromolecular densities (8-40% cytoplasmic occupancy). Controls were carried out in buffers without crowders. Protein structure and stability under crowding were monitored by circular dichroism (CD) spectroscopy (Supplementary Information).

### Thiol-labeling experiments

Cysteine solvent accessibility was measured using 5,5′-dithiobis-(2-nitrobenzoic acid) (DTNB, Ellman’s reagent, Sigma). Labelling reactions were initiated by mixing 10 µM of protein with DTNB at the indicated concentrations in labeling buffer (20 mM Tris, pH 8.0, 150 mM NaCl) at 25 °C. Release of TNB^2-^ was monitored by absorbance at 412 nm until a steady state (∼300 s) was reached using a stopped-flow spectrophotometer (Applied Photophysics).

Individual kinetic traces were baseline-corrected against buffer controls and globally fit to as many exponential terms as cysteines present in the construct, yielding observed rate constants 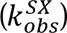 for each site at each DTNB concentration. Fits were performed in Python (SciPy), and reported values represent the mean ± SD from at least three independent measurements. The dependence of 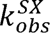 on DTNB concentration was modeled using the Linderstrøm-Lang model as described previously.^40^

### HDX-MS experiments

Protein samples (∼150 µM) were diluted 10-fold into freshly prepared deuteration buffer (10 mM HEPES, 150 mM NaCl, 1 mM MgCl2, 2mM TCEP adjusted to pD 7.0 with NaOD) in 99% D2O at 25 °C. Exchange was carried out for labeling times ranging from 20 s to 24 h. At each time point, aliquots were mixed with quench buffer (120 mM glycine-HCl, 150 mM NaCl, 2 mM TCEP, adjusted to pH 2.5 with HCl, ∼0 °C) to halt labelling. Quenched samples were immediately processed for digestion.

Proteins were digested online using an immobilized pepsin column maintained at 0 °C, and the resulting peptides were captured on a TARGA C8 5 µm Piccolo HPLC column (1.0 × 5.0 mm; Higgins Analytical) for desalting. Peptides were then separated on an analytical TARGA C8 HPLC column (0.3 × 75 mm; Higgins Analytical) using a linear 10-40% acetonitrile gradient (Buffer A: 0.1% formic acid; Buffer B: 0.1% formic acid in 99.9% acetonitrile) at 8 µL/min, with all chromatography conducted at 0 °C.

Eluting peptides were analyzed on a QExactive mass spectrometer (Thermo Fisher Scientific) equipped with electrospray ionization. Spectra were collected over an m/z range of 200-2000. Undeuterated controls were analyzed in data-dependent mode for peptide identification, and peptide lists were curated in Proteome Discoverer (Thermo Fisher Scientific).

Deuterium uptake was quantified by centroid analysis of isotopic envelopes using EXMS2.^41^ Percent uptake was normalized to a fully deuterated control (24 h in D O at 25 °C, followed by heating at 45 °C for 15 min) using the following equation:

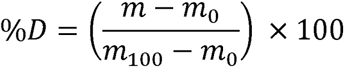

where *m, m_0_*and *m_100_* are the observed centroid masses of the peptide at a given time point, the non-deuterated peptide and fully deuterated peptide, respectively.^42^

Uptake curves were fit to single, double, or triple exponential models in Python (SciPy), and the observed exchange rates 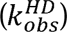 are reported as mean ± SD from replicate experiments. Intrinsic exchange rates (*k_int_*) were calculated as reported previously,^43, 44^ and protection factors (PF) were derived as 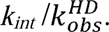

## Acknowledgements

We thank the Johnson Foundation Structural Biology and Biophysics Core at the University of Pennsylvania for HDX-MS resources. This work was supported by the National Science Foundation through NSF MCB 2218156 (G.R.B.).

## Data Availability Statement

The data that support this study are available from the corresponding author upon reasonable request. The HDX-MS data have been deposited to the PRIDE database. Referenced structures are: PDB ID 3FKE and 3L25.

