## Supplementary Information for "Opening of a cryptic pocket is not suppressed by excluded volume effects of crowding"

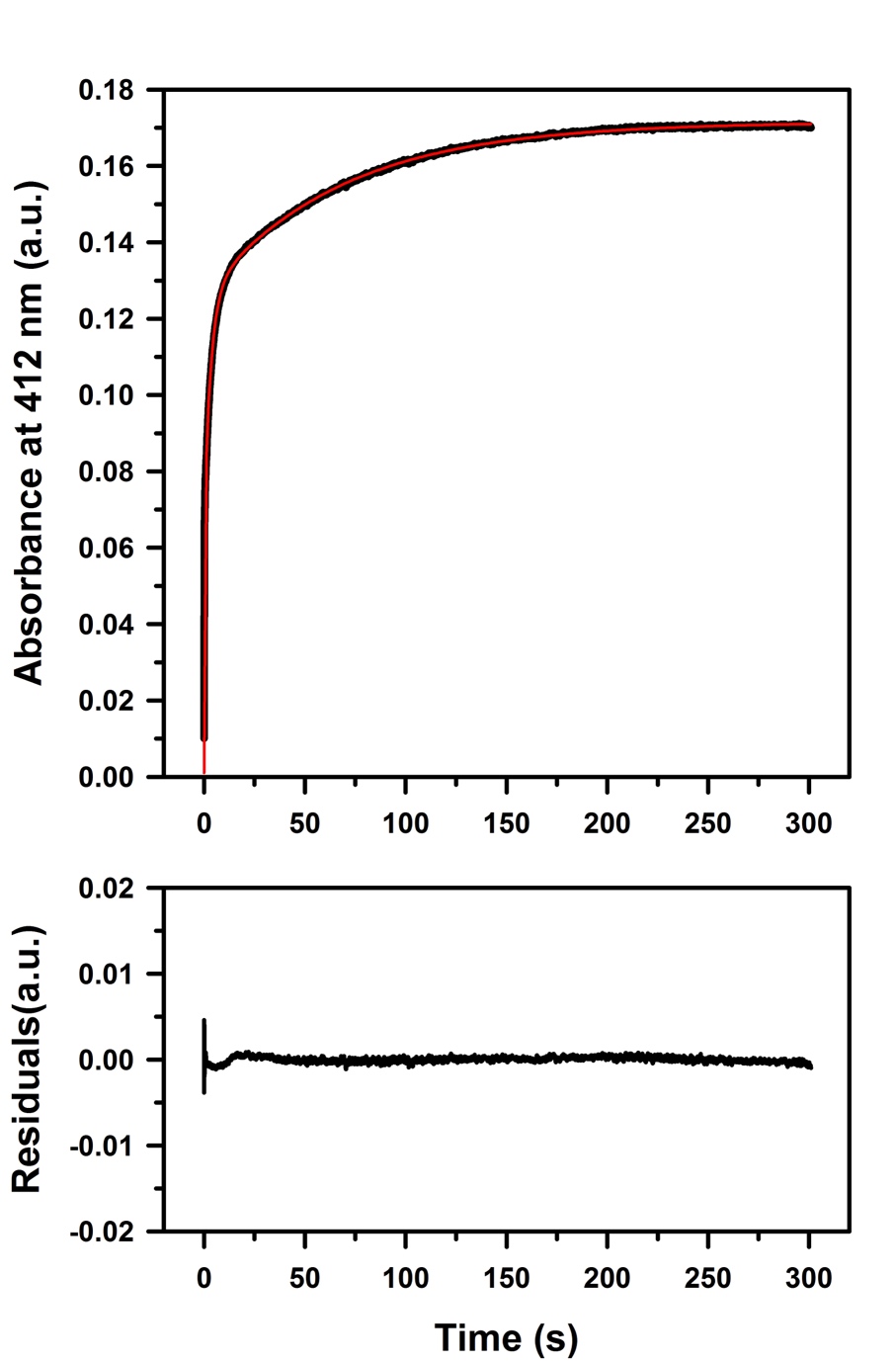

**Supplementary Figure 1:** Representative thiol-labeling time trace and second-exponential fit for the VP35 C247S/C275S variant.

A representative thiol-labeling time trace (black, top) collected at 125 µM DTNB and 5 µM protein is shown with the corresponding double-exponential fit (red). The data were background-subtracted using the average signal from DTNB-only control reactions to correct for spontaneous DTNB hydrolysis. The residuals (bottom) are centered around zero with no systematic deviations, indicating that the fit appropriately captures the kinetics.

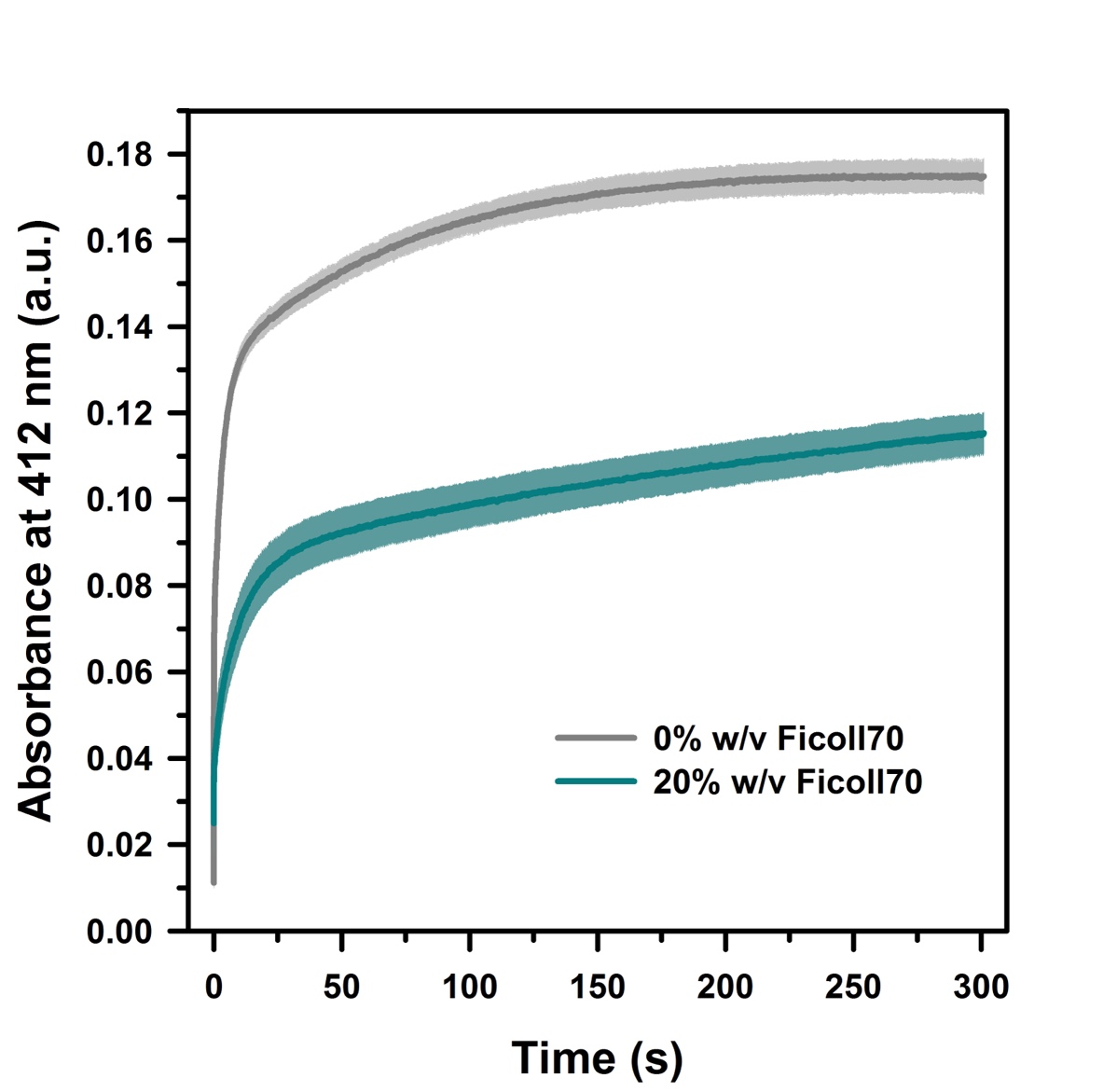

**Supplementary Figure 2:** Representative thiol-labeling kinetics of VP35 C247S/C275S variant under dilute and 20% w/v Ficoll 70 conditions.

Representative buffer-subtracted absorbance traces at 412 nm for 5 μM protein labeled with 125 μM DTNB in buffer (0% w/v Ficoll 70, gray) and in the presence of 20% w/v Ficoll 70 (teal). Traces represent the mean of three independent measurements, and shaded regions indicate standard deviations.

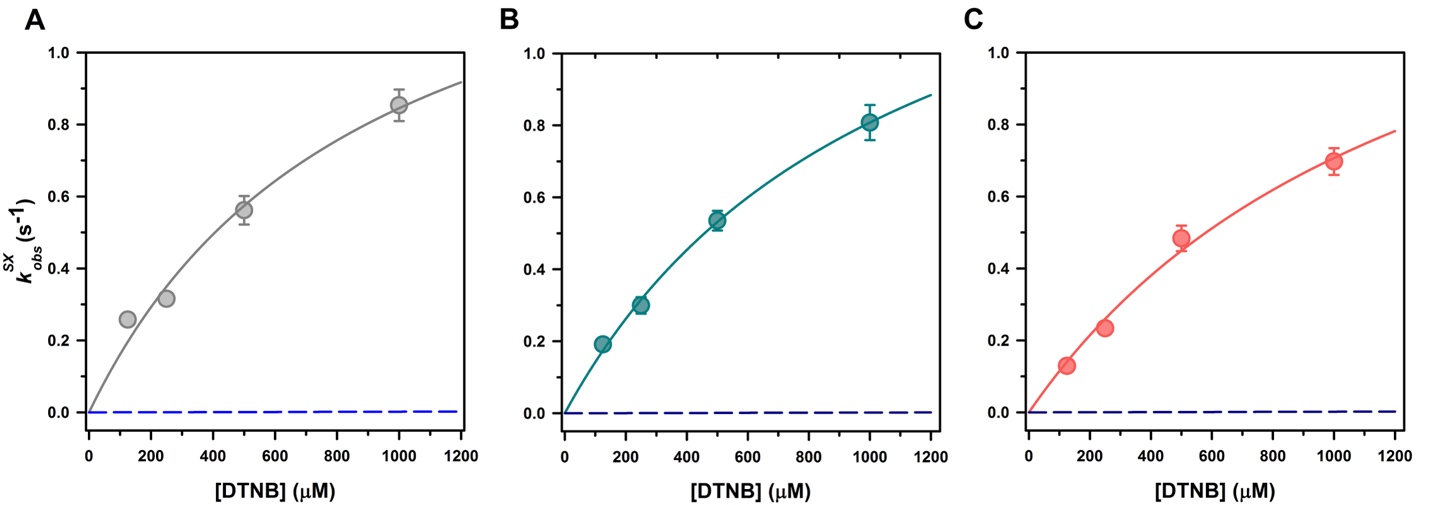

**Supplementary Figure 3:** Determination of cryptic pocket opening from C307 thiol-labeling kinetics under Ficoll 70 crowding conditions.

(A–C) Dependence of the observed labeling rate ($k_{\mathrm{obs}}^{\mathrm{SX}}$) for C307 on DTNB concentrations in the C247S/C275S VP35 IID background in the presence of 0 mg/mL (A), 100 mg/mL (B), and 200 mg/mL Ficoll 70 (C). Symbols represent experimentally measured rate constants obtained from thiol-labeling experiments, and solid lines show fits to the EXX thiol-exchange model. Error bars represent standard deviations from independent measurements. The blue dashed line shows the predicted labeling rate assuming exchange occurs through the unfolded state.

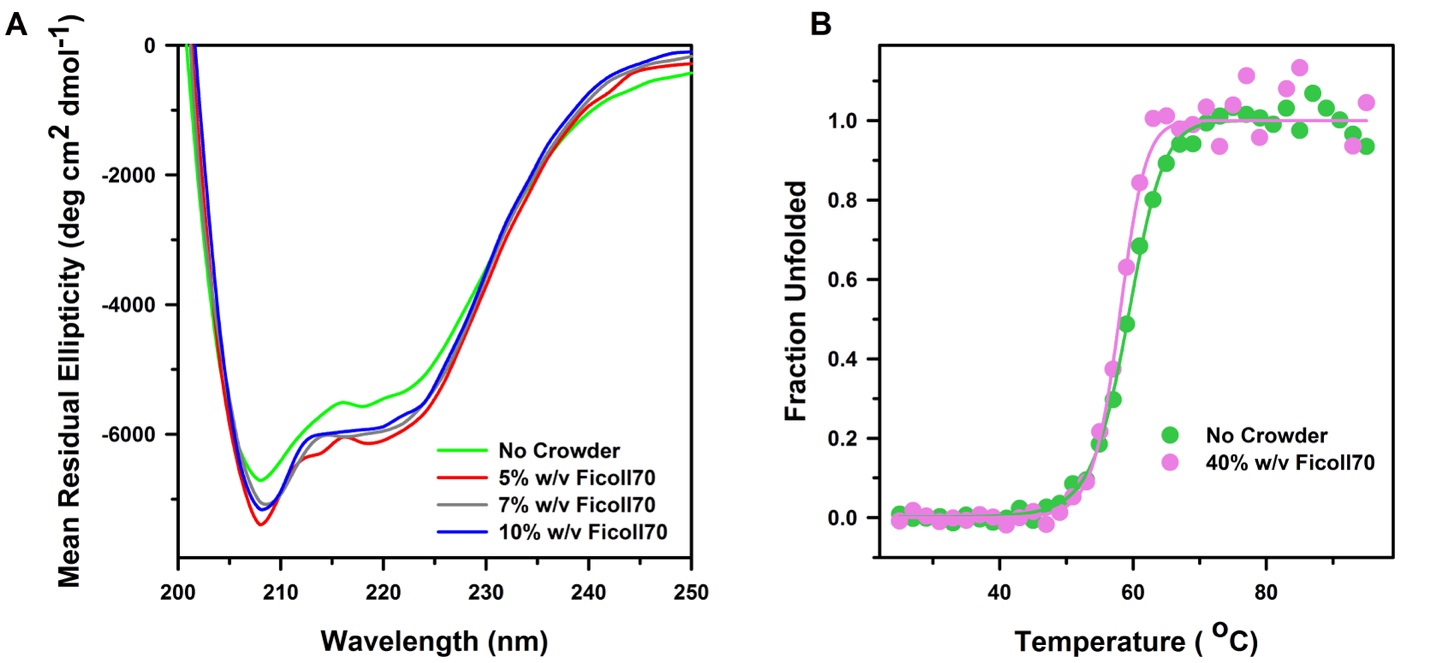

**Supplementary Figure 4:** Secondary structure and thermal stability of VP35 IID in the presence of Ficoll 70.

**(A)** Far-UV circular dichroism (CD) spectra of VP35 IID collected in the absence of crowder and in the presence of 5%, 7%, and 10% w/v Ficoll 70. **(B)** Thermal denaturation of VP35 IID monitored by circular dichroism in the absence and presence of 40% w/v Ficoll 70. Fraction unfolded was calculated by normalizing the CD signal between the folded and unfolded baselines and plotted as a function of temperature. Solid lines represent fits to a two-state unfolding model.

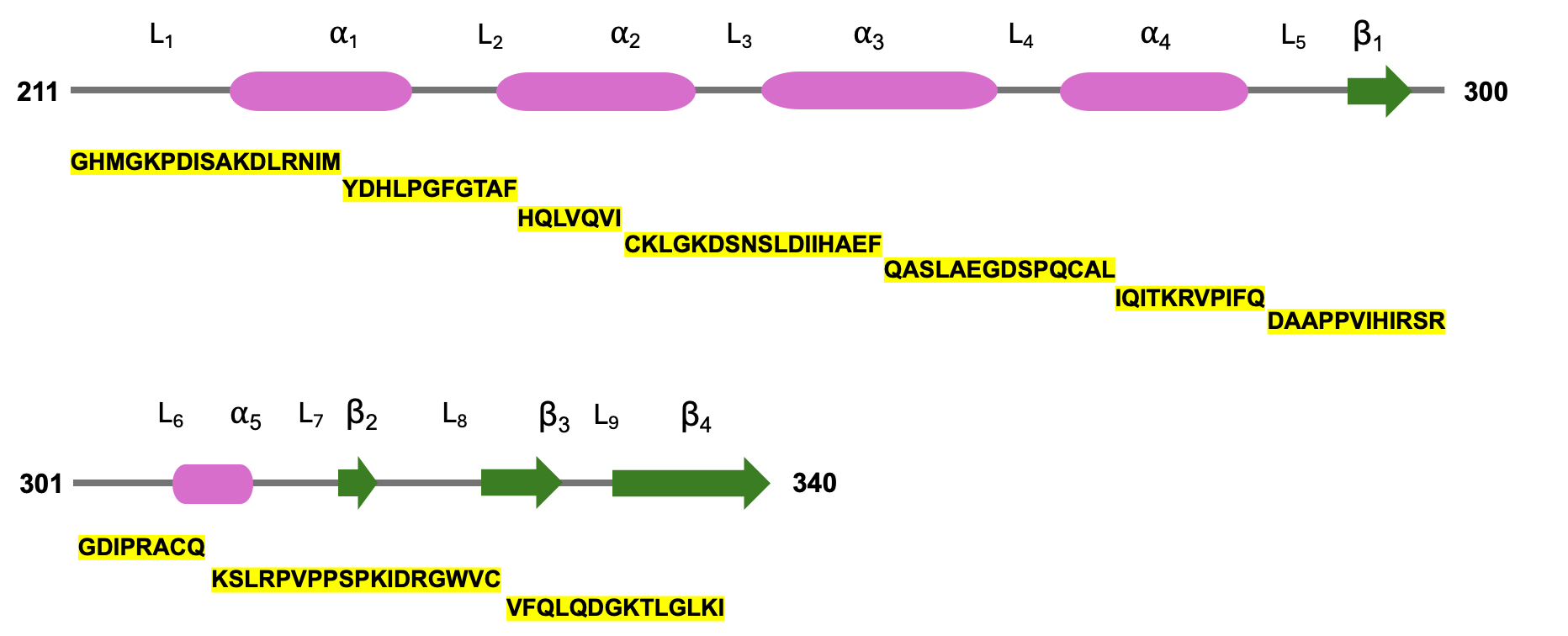

**Supplementary Figure 5: HDX-MS peptide coverage across the VP35 IID sequence and secondary structure map.**

The VP35 IID domain is shown with annotated secondary structure elements (helices in magenta, loops in gray, and β-strands in green). Peptides detected by HDX-MS are highlighted in yellow along the linear sequence. Coverage spans over the whole protein, including regions that form the RNA-binding interface and the cryptic pocket. This peptide resolution enables mapping of local protection changes and supports structural interpretation of HDX-MS results.

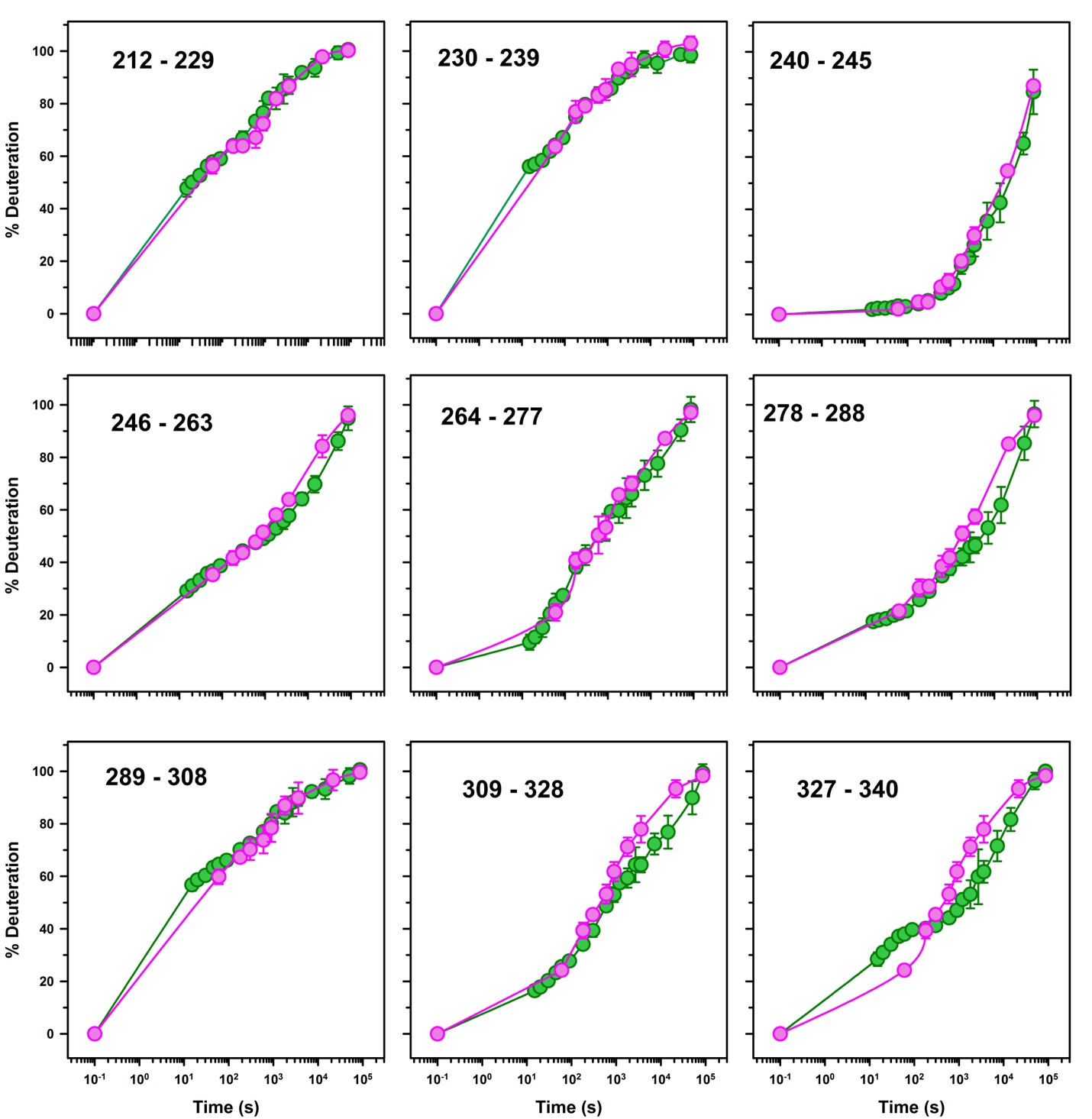

**Supplementary Figure 6: HDX-MS uptake kinetics across WT VP35 IID peptides under dilute and Ficoll 70 crowded conditions.**

Percent deuterium uptake as a function of labeling time is shown for representative peptides spanning the VP35 IID sequence. Data collected in buffer (green) and in 40% w/v Ficoll 70 (magenta) are shown for each peptide. Data points represent the mean of at least three independent measurements, and error bars indicate standard deviations.

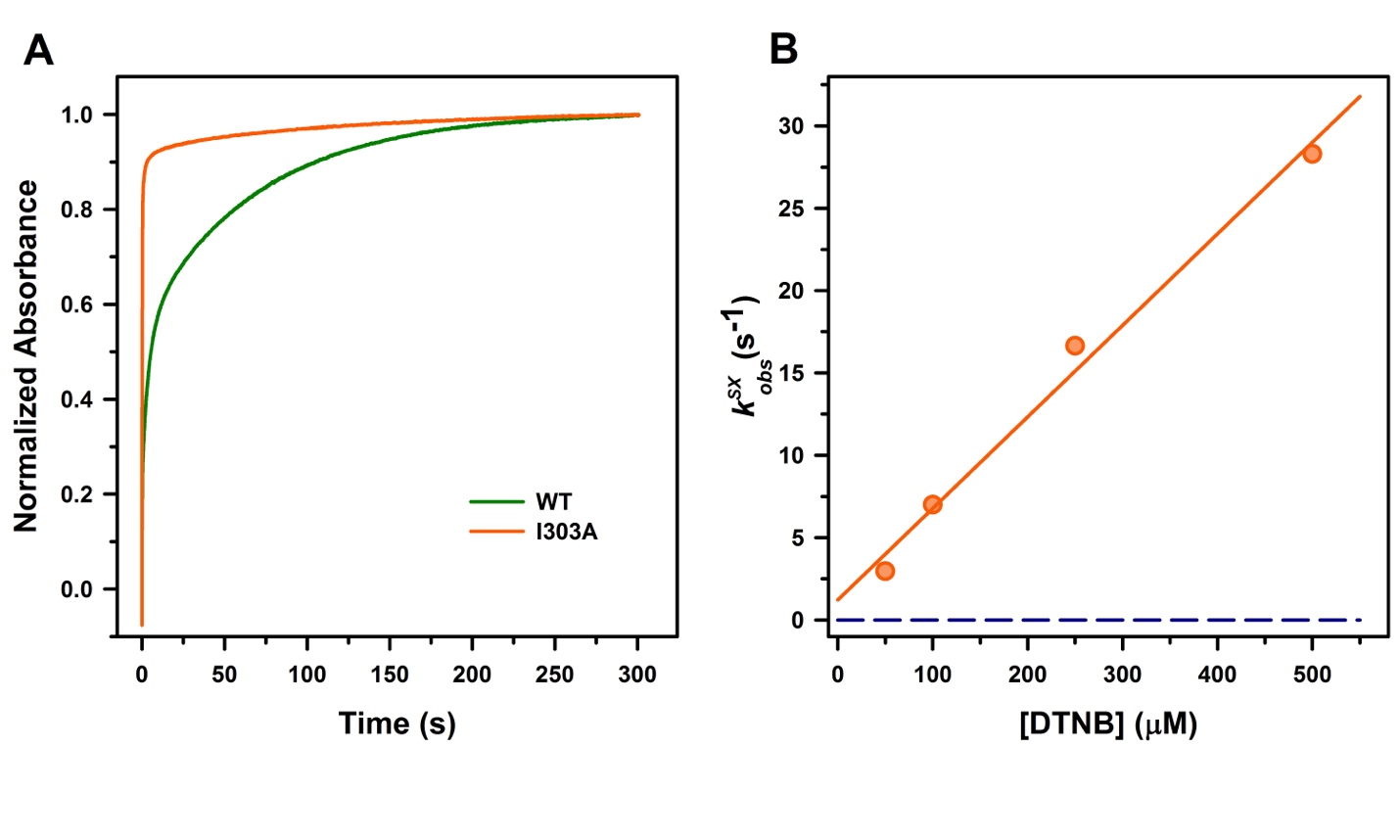

**Supplementary Figure 7:** Increased cryptic pocket accessibility in the I303A variant as measured by thiol exchange experiments.

(A) Representative thiol-labeling traces for WT VP35 IID and the open-state stabilized variant I303A, measured at 125 μM DTNB. Absorbance values were normalized to the final signal to facilitate comparison of labeling kinetics. (B) Dependence of the observed labeling rate ($k_{\mathrm{obs}}^{\mathrm{SX}}$) for C307 in the I303A variant on DTNB concentration. The solid line shows the fit to the EX2 thiol-exchange model. Error bars represent standard deviations from three independent measurements. The blue dashed line indicates the expected labeling rate from the unfolded state.

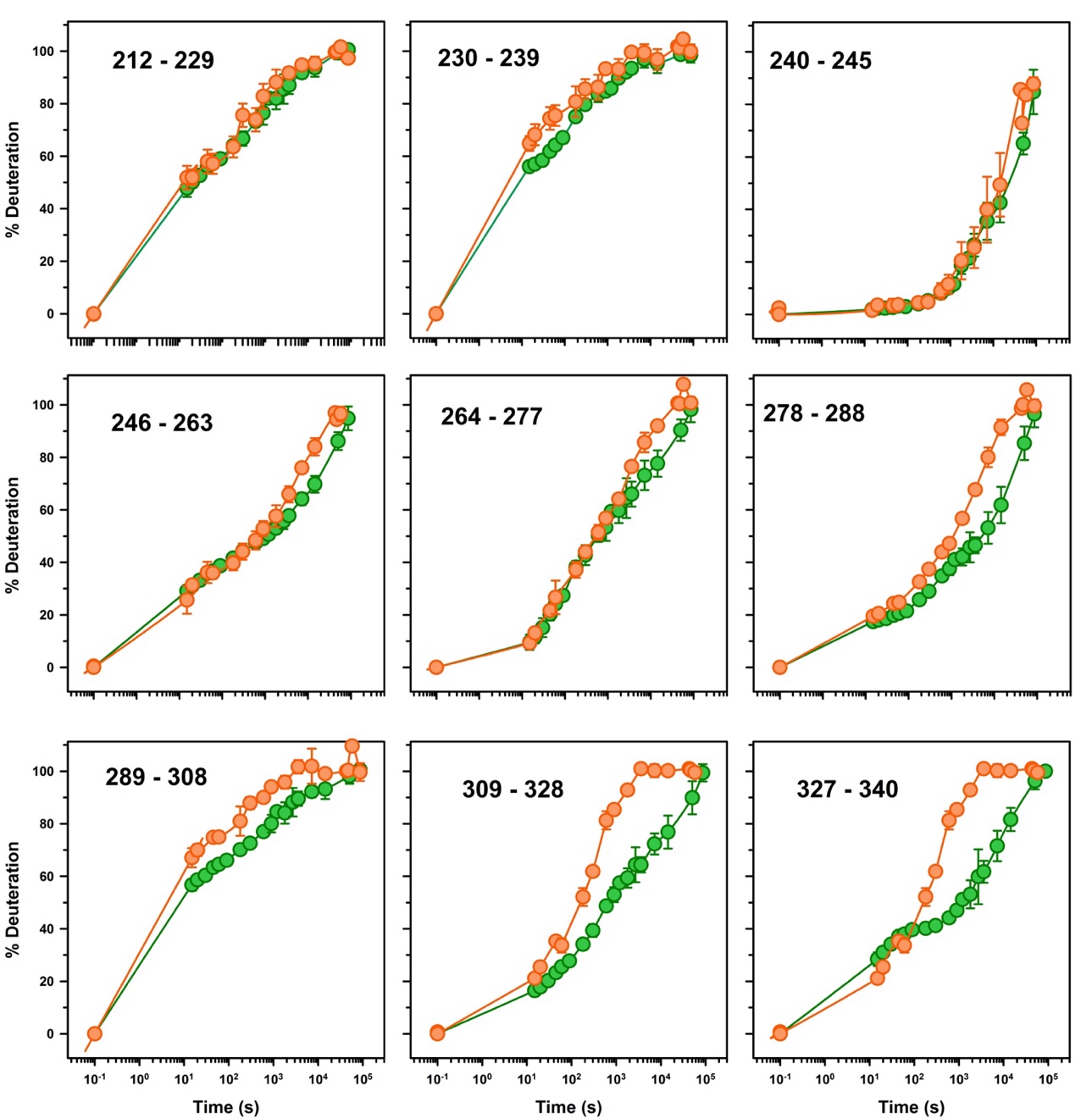

**Supplementary Figure 8:** HDX-MS uptake kinetics of WT and I303A VP35 IID under dilute conditions.

Percent deuterium uptake as a function of labeling time for all peptides spanning the VP35 IID sequence. Data collected for WT VP35 IID (green) and the I303A variant (orange) are shown for each peptide. Increased deuterium uptake is localized primarily to peptides spanning the cryptic pocket, whereas peptides outside the pocket region exhibit minimal differences between WT and I303A. Data points represent the mean of at least three independent measurements, and error bars indicate standard deviations.

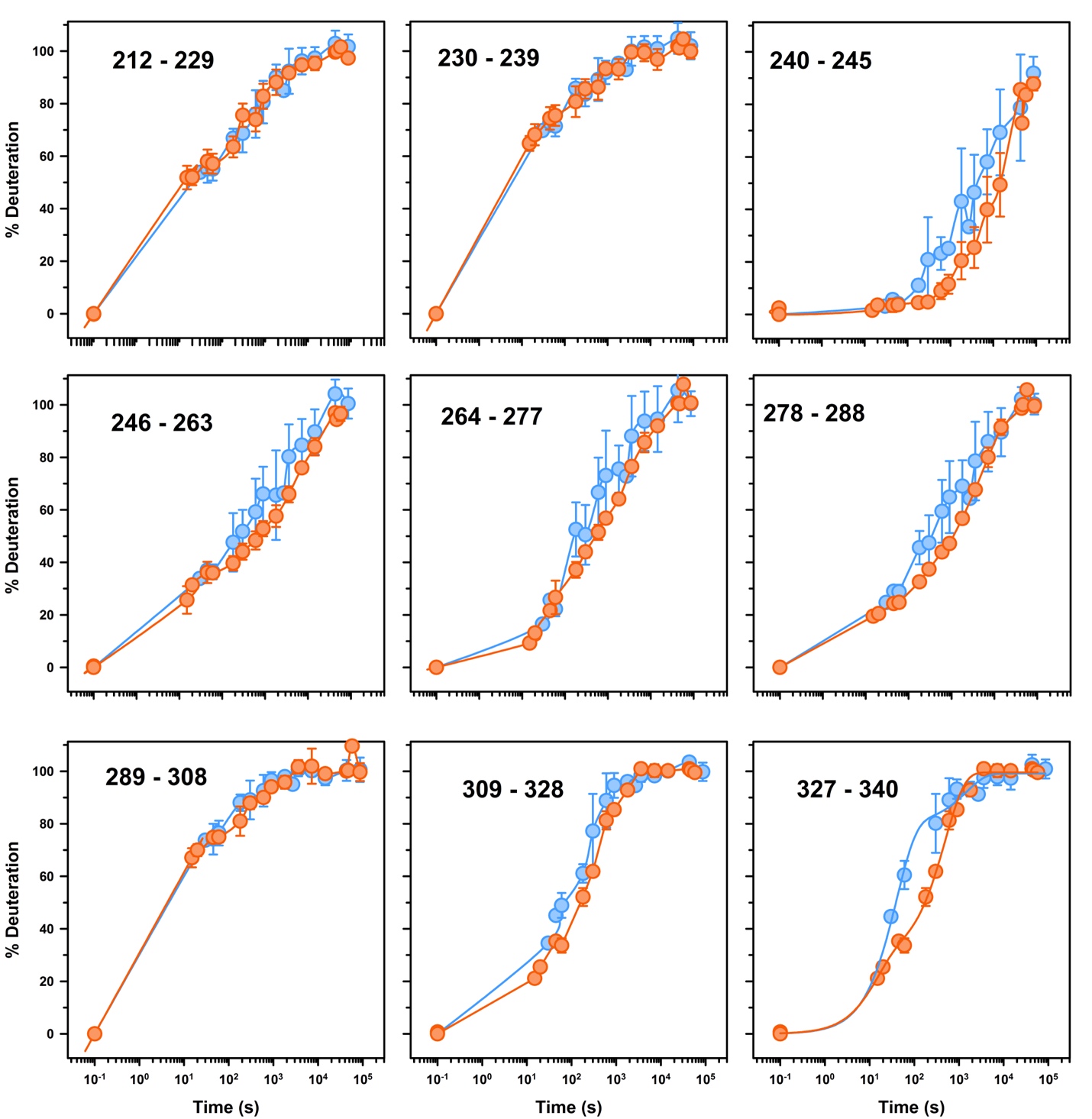

**Supplementary Figure 9:** HDX-MS uptake kinetics of I303A VP35 IID under dilute and Ficoll 70 crowded conditions.

Percent deuterium uptake as a function of labeling time for all peptides spanning the I303A VP35 IID sequence. Data collected for the I303A variant in buffer (orange) and in the presence of 40% w/v Ficoll 70 (blue) are shown for each peptide. Representative peptides from the cryptic pocket region (D289–Q308 and K309–F328) are presented in Figure 5. Data points represent the mean of three independent measurements, and error bars indicate standard deviations.

**Supplemental Methods:**

**Far-UV Circular Dichroism Spectroscopy:**

Far-UV circular dichroism (CD) spectra were collected to assess secondary structure integrity of VP35 IID under crowding conditions. Measurements were performed on a JASCO J-1500 spectrophotometer equipped with a Peltier temperature controller. Protein samples (WT VP35 IID) were prepared at 15 µM in 20 mM HEPES pH 7.4, 150 mM NaCl and 2 mM TCEP. Ficoll 70 was added at the indicated weight/volume concentrations (0 - 40% w/v), and the samples were equilibrated for 30 min at 25 °C prior to data acquisition.

CD spectra from 200 - 250 nm were recorded at 25 °C using a 0.1 cm quartz cuvette, 1 nm bandwidth, 2 nm step size, and 1s averaging time. Each spectrum represents the average of three accumulations. Buffer baselines containing the corresponding Ficoll 70 concentration were collected in parallel and subtracted from the sample spectra. Mean residue ellipticity was calculated using standard procedures.

**Thermal denaturation Experiments:**

Thermal denaturation was monitored at 222 nm from 25 - 95 °C at a heating rate of 1 °C/min. Samples contained either no crowder or 40% w/v of Ficoll 70. Raw ellipticity values at 222 nm were baseline-corrected and converted to fraction unfolded (FU) using a standard two-state model:

$$FU= \frac{\theta_{T}-\theta_{N}}{\theta_{U}-\theta_{N}}$$

where $\theta_{T}$ is the measured ellipticity at temperature T, and $\theta_{N}$ and $\theta_{U}$are the signals of native and unfolded baselines.
